# Development of a high-throughput γ-H2AX assay based on imaging flow cytometry

**DOI:** 10.1101/637371

**Authors:** Younghyun Lee, Qi Wang, Igor Shuryak, David J. Brenner, Helen C. Turner

**Affiliations:** Center for Radiological Research, Department of Radiation Oncology, Columbia University Irving Medical Center, New York, NY 10032, USA

**Keywords:** imaging flow cytometry, DNA repair kinetics, human lymphocytes, high throughput, radiation sensitivity, ionizing radiation

## Abstract

**Background:** Measurement of γ-H2AX foci formation in cells provides a sensitive and reliable method for quantitation of the radiation-induced DNA damage response. The objective of the present study was to develop a rapid, high-throughput γ-H2AX assay based on imaging flow cytometry (IFC) using the ImageStream^®X^ Mk II (ISX MKII) platform to evaluate DNA double strand break (DSB) repair kinetics in human peripheral blood cells after exposure to ionizing irradiation.

**Methods:** The γ-H2AX protocol was optimized for small volumes (100 µl) of blood in Matrix™ 96-tube format and blood cell lymphocytes were identified and captured by ISX INSPIRE™ software and analyzed by Data Exploration and Analysis Software.

**Results:** Presented here are: 1) dose response curves based on γ-H2AX fluorescence intensity and foci number, 2) measurements of DNA repair kinetics up to 24 h after exposure to 4 Gy γ rays and, 3) a mathematical approach for modeling DNA DSB rejoining kinetics using two key parameters a) rate of γ-H2AX decay, and b) yield of residual unrepaired breaks.

**Conclusions:** The results indicate that the IFC-based γ-H2AX protocol may provide a practical, high-throughput and inexpensive platform for measurements of individual global DSB repair capacity and facilitate the prediction of precision medicine concepts.

## Background

Double Strand Breaks (DSBs) are one of the most important types of DNA damage. DSBs are more difficult to repair than many other lesions and their incorrect repair (e.g., misrejoining of broken DNA strands from different chromosomes) can result in cytotoxic or carcinogenic genomic alterations. Defects in the DNA repair machinery may increase cell vulnerability to DNA-damaging agents and accumulation of mutations in the genome, and could lead to the development of various disorders including cancers. Epidemiological evidence supports a strong association between global DSB repair capacity and cancer risk (1–3), radiation sensitivity (4, 5) and response to cancer therapy (6, 7). The association between genetic defects in DNA repair and increased clinical radiosensitivity has been identified in many studies and used as a basis for the development of predictive assays for normal tissue toxicity (8).

Over the past decade, the γ-H2AX assay has been applied to a variety of cell types and tissues to correlate γ-H2AX levels with DNA damage and repair (9–13). Following radiation exposure, histone H2AX is rapidly phosphorylated by ATM and/or DNA-PK kinases at or near the vicinity of DNA DSB sites to form γ-H2AX (14). Immunolabeling of γ-H2AX provides a quantitative measurement and direct visualization of DSBs as fluorescent nuclear foci. At the cellular level, the kinetics of formation or loss of γ-H2AX foci may reflect the rate or efficiency of DSB repair (15). The biphasic nature of DSB repair kinetics has been associated with different repair pathways that allow repair for a fast (initial few hours) and slow component (hours to days) of repair (16, 17). Additionally, there is evidence that the DSBs assayed several hours after the initial radiation challenge that still remain unrepaired known as residual DNA damage, may be predictive of individual susceptibility to complex DNA lesions that can be lethal (18). Current evidence suggests that there is a large inter-individual variation in DSB DNA repair capacity in lymphocytes from healthy individuals (19–21). Further, clinical radiosensitivity is often linked to defects in DNA repair (5, 22, 23). The capacity to repair DSB is therefore an important factor to consider in risk assessment, however studies to date are limited due to no large-scale prospective evidence or ability to conduct high-throughput phenotypic assays (24).

The objective of the present study was to develop a rapid, high-throughput γ-H2AX assay based on imaging flow cytometry (IFC) using the ImageStream^®X^ Mk II (ISX MKII) platform to evaluate DNA DSB repair kinetics in human peripheral blood cells after exposure to ionizing irradiation. Imaging flow cytometry is a relatively new technique which combines the speed of flow cytometry with the imaging capability of conventional microscopy (25–27). It has been used to analyze cell death, apoptosis and immune response as an advanced method for fluorescence-based analysis of cellular morphology and heterogeneity (28–33). Combining the strength of flow cytometry and conventional microscopy enables high throughput characterization of cells on a microscopic scale(34). This approach was applied to develop high throughput γ-H2AX assay. We demonstrate this technique here and present: 1) dose response curves based on γ-H2AX fluorescence intensity and foci number, 2) measurements of DNA repair kinetics up to 24 h after exposure to 4 Gy γ rays and, 3) a mathematical approach for modeling DSB rejoining kinetics using two key parameters a) rate of γ-H2AX decay, and b) yield of residual unrepaired breaks.

## Methods

### Blood collection and irradiation

Blood was collected by venipuncture in 5 mL lithium-heparinized Vacutainer^®^ tubes (BD Vacutainer™, Franklin Lakes, NJ) from healthy adult donors (2 female and 2 male) with informed consent and approval by the Columbia University Medical Center Institutional Review Board (IRB protocol IRB-AAAE-2671) in 5 mL lithium-heparinized Vacutainer^®^ tubes (BD Vacutainer™, Franklin Lakes, NJ). All donors were non-smokers in relatively good health at the time of donation with no obvious illnesses such as colds, flu, or infections and no known exposures to medical ionizing radiation within the last 12 months. Fresh blood aliquots (1 mL) were dispensed into 15 mL conical bottom tubes (Santa Cruz Biotechnology, Dallas, TX) and were irradiated with γ rays (0, and 4 Gy) using a Gammacell^®^ 40 ^137^Cesium irradiator (Atomic Energy of Canada, Ltd., Chalk River, ON). The blood sample tubes were placed on their side in the middle of the chamber and irradiated with a dose rate of 0.73 Gy/min (35). The ^137^Cs irradiator was calibrated annually with TLDs and homogeneity of exposure across the sample volume was verified using EBT3 Gafchromic film with less than 2% variation within the sample (Ashland Advanced Materials, Gafchromic, Bridgewater, NJ).

### γ-H2AX assay immunolabeling protocol

Immediately after irradiation, 100 μl blood aliquots transferred to 1.4 mL 2D Matrix™ microtubes (Thermo Scientific™, Waltham, MA) containing 900 µL RPMI 1640 culture medium (Gibco, Waltham, MA) supplemented with 15% FBS and 2% Penicillin and Streptomycin (all reagents from Invitrogen, Eugene, OR). The rack containing microtubes was placed into an incubator at 37°C, 5% CO_2_ up to 24 h. At specific time points after irradiation (0.5, 1, 3, 6 and, 24 h), cultured blood samples were lysed and fixed with 1X Lyse/fix solution (BD Phosflow™,; BD Biosciences,, San Jose, CA), washed with 1X phosphate buffered saline (PBS, Gibco, Gaithersburg, MD), suspended in 50% cold methanol, and stored at 4 °C for 24 h. Fixed cells were permeabilized with 0.1% Triton X-100 (Sigma-Aldrich, St. Louis, MO) at room temperature for 10 min and then incubated with Alexa Fluor^®^ 488 Mouse anti-H2AX (pS139) antibody (clone N1-431, BD Pharmingen™, Franklin Lakes, NJ), diluted 1:1000 with 1% bovine serum albumin (BSA, Sigma-Aldrich, St. Louis, MO) at 4°C overnight, after which the samples were washed with 1X PBS and stained with 5 μM DRAQ5™ (Thermo Scientific™) at RT for a minimum of 5 min. All solution transferring or mixing in microtubes was performed using a 1.2-ml multichannel electronic pipet (Eppendorf Inc., Westbury, NY). All steps in the procedure were performed at room temperature (RT) and microtubes in racks were spun at 250×g for 3 min.

### Data acquisition and analysis on the ISX and IDEAS^®^

The 96-well plate of samples were transferred to the ImageStream^®X^ Mk II (ISX MKII) imaging flow cytometer (LUMINEX Corporation, Austin, Texas) for automated sample acquisition and captured using the ISX INSPIRE™ data acquisition software. Images of 5000~12,000 cells were acquired at 40X magnification using the 488 nm excitation laser at 200 mW: Bright field (BF) images were captured on channel 1, γ-H2AX immunostaining on channel 2, DRAQ5 images on channel 5 and side scatter on channel 6. Data was collected with only the Area feature applied, such that events with areas less than 60 pixels (15 μm^2^) were gated out in order to minimize the collection of small debris. For the compensation, cells stained with γ-H2AX antibody or DRAQ5 only and were captured using the 488 nm laser without brightfield illumination. The compensation coefficients were acquired automatically by the IDEAS 6.2 compensation wizard. To quantify the γ-H2AX expression levels, the viable lymphocytes population was gated for foci quantification and total γ-H2AX fluorescence intensity. Nuclear foci formation was identified using the wizard in IDEAS which automated targeting and enumerate the foci. The geometric mean of the γ-H2AX fluorescence intensity of individual cells from each sample was analyzed. For the dose response curve, γ-H2AX foci and intensity levels were measured at 1 h post irradiation. All curves were generated using GraphPad Prism 7 (GraphPad software Inc., La Jolla, CA), and R^2^ value was calculated to assess goodness of fit of curves from linear regression analysis.

### Quantitative modeling of DNA repair kinetics

For the kinetic curves, γ-H2AX levels were measured at 0.5, 1, 3, 6 and, 24 h after 4 Gy irradiation. The data on γ-H2AX foci (F) at different time points (T) after irradiation were quantitatively modeled by the following equation, where F_bac_ is the background value prior to irradiation, F_res_ is the residual value remaining at long times (e.g. 24 h) after irradiation, K_prod_ is the constant for induction of foci by radiation, and K_dec_ is the constant for decay of foci after irradiation (20):

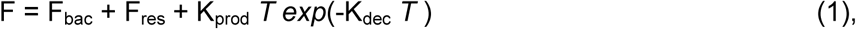

We used least squares fitting in Maple 2017 software (https://www.maplesoft.com/) as a practical approach for estimating K_dec_ and F_res_, involving curve fitting of each sample data set to Eq. (1). Thus, as we propose below we will use both the decay constant (K_dec_) and residual excess fluorescence intensity (F_res_) to describe each individual’s DNA DSB Repair Capacity.

## Results

### Development of IFC-based high throughput γ-H2AX assay

We have developed a simple and rapid IFC-based γ-H2AX protocol which comprises of the following four components: (1) Sample preparation of finger-stick sized blood samples (< 100 µl) in 96 well format, 2) Automated cellular image acquisition of immunofluorescent-labelled biomarkers using the ISX MKII system (3) Quantification of γ-H2AX biomarker levels using Image Data Exploration and Analysis Software and, (4) Quantitative modeling of DNA repair kinetics in peripheral blood lymphocytes. Figure 1 shows schematic work flow for the IFC-based γ-H2AX protocol. In general, the immunolabeling protocol is less than 2 hours while the acquisition and analysis of each sample (~3000 cells) can be finalized within 3 minutes.

**Figure 1.**
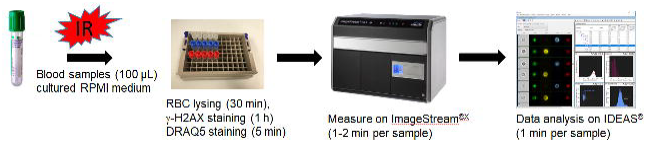
Development of a simple and fast γ-H2AX assay protocol. Fresh blood samples (100 µL) were prepared and cultured with RPMI medium after gamma irradiation. At specific time points up to 24 h after irradiation, whole blood samples were lysed, fixed and stained with γ-H2AX antibody and the nuclei counter-stained with DRAQ5. Imagery of cells were collected by the ImageStream^®X^ (ISX) Mark II imaging flow cytometer for automated sample acquisition and captured using the ISX INSPIRE™ software. The acquired data was analyzed by IDEAS® software.

### Quantification of γ-H2AX levels using IDEAS software

Figure 2 shows the gating strategy to identify γ-H2AX levels in non-apoptotic human lymphocytes from the cell population. The focused cells were gated according to the gradient similarity feature by visual inspection of cell images in the brightfield channel (Fig. 2A). Single cells were then selected from images according to their area and aspect ratio in the brightfield channel (Fig 2B) and nucleated cells are selected based on DRAQ5 positivity to exclude the dead cells (Fig 2C). Given that the level of γ-H2AX in granulocytes is barely affected by radiation (36), lymphocytes are gated according to their area on bright field and side scatter for further measurement of the γ-H2AX fluorescence intensity and foci formation (Figure 2D). Pan-nuclear γ-H2AX stained cells displayed a typical apoptotic pattern (Figure 3A) and were increased with the time post irradiation (Figure 3B), thus were excluded from the γ-H2AX analysis. For each data point, 8273 ± 317 cells were analyzed from 100 µl whole blood within 1 – 2 min. Gamma H2AX yields were measured in 2076 ± 123 (mean ± SEM) non-apoptotic lymphocytes in average.

**Figure 2.**
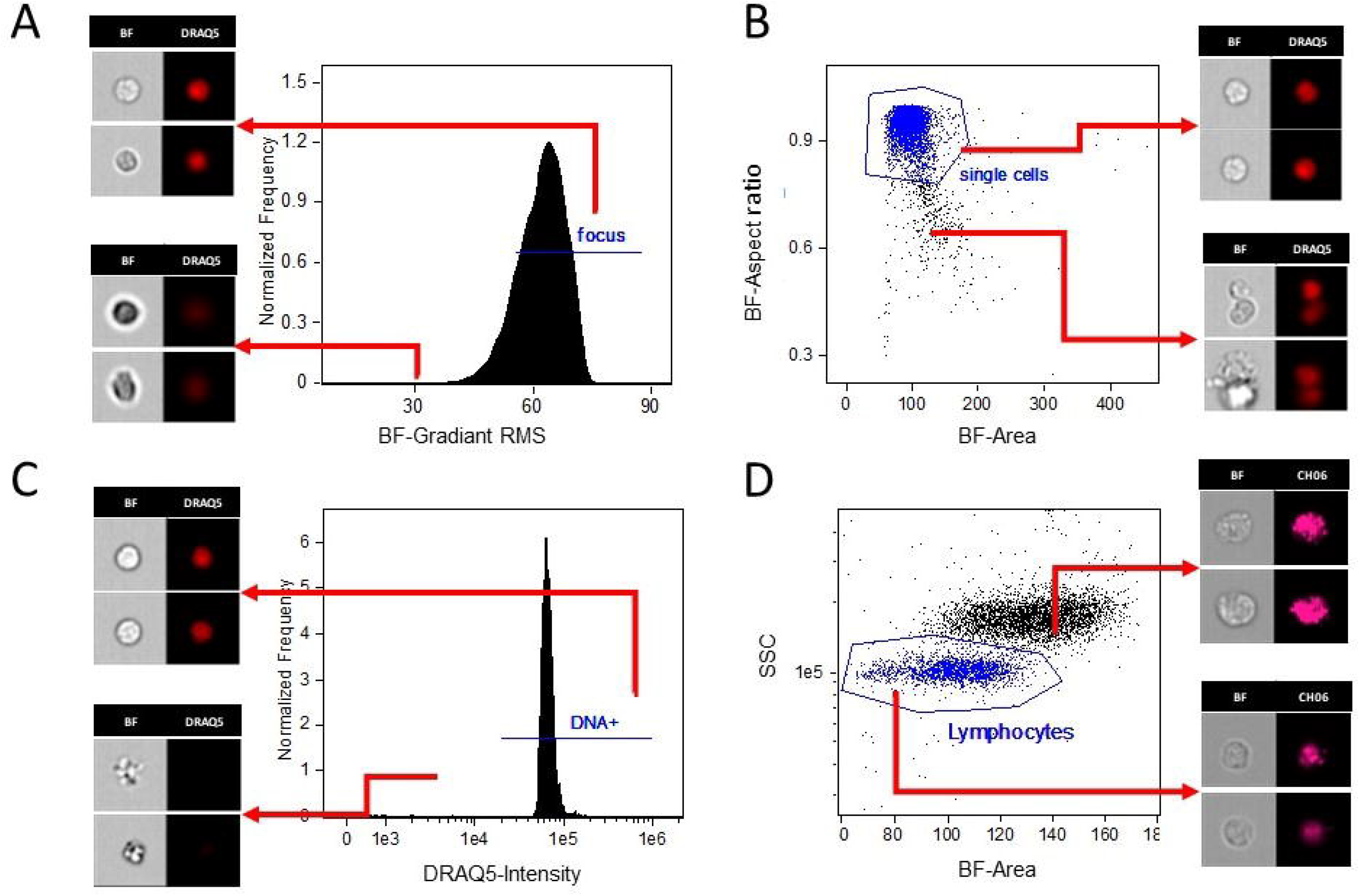
Gating strategy for assessing γ-H2AX levels in the IDEAS^®^ software. (A) According to the gradient similarity feature in the brightfield channel, the focused cells were gated. (B) According to their area and aspect ratio in the brightfield channel, single cells were then selected from images. (C) Nucleated cells are selected based on DRAQ5 positivity. (D) According to cell area on bright field and side scatter, lymphocytes are selected for γ-H2AX analysis. BF=Bright field.

**Figure 3.**
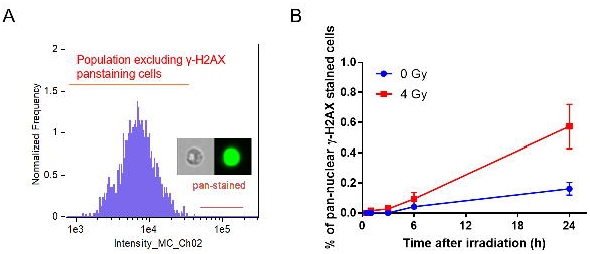
Percentages of pan-nuclear γ-H2AX stained cells increase with time in irradiated and non-irradiated cells. (A) Gating of pan-nuclear γ-H2AX stained cells. (B). Percentages of pan-nuclear γ-H2AX stained cells. The data presented as mean ± SEM.

The mean fluorescence intensity of γ-H2AX within nuclear boundary of individual cells was analyzed and exported from the IDEAS® software. The number of γ-H2AX foci was calculated using the spot counting wizard in IDEAS software as shown in Figure 4. The spot counting wizard automatically creates masks based on calculation of more than selected 30 low foci cells and 30 high foci cells by visual speculation. The mask is composed of three different masks on channel 2 and channel 5 (i) Spot mask identified spots with a size<1 pixel and a spot to background ratio>4.5; (ii) Peak mask identified intensity areas from an image with local maxima (bright) or minima (dark); (iii) Range identified spot image with a size<200 pixel and aspect ratio from 0 to 1; (iv) Overlapping with Channel 5. The representative foci mask is shown in Figure 4. Finally, the feature Spot count was used to enumerate foci identified by the mask. A template file was then generated and applied to all samples using a batch analysis. Using this imaging flow cytometry-based system, dose- and time-dependent γ-H2AX levels responding to radiation exposure were measured automatically over 24 h yielding an estimate of global DSB repair capacity as well as a measure of unrepaired DSB.

**Figure 4.**
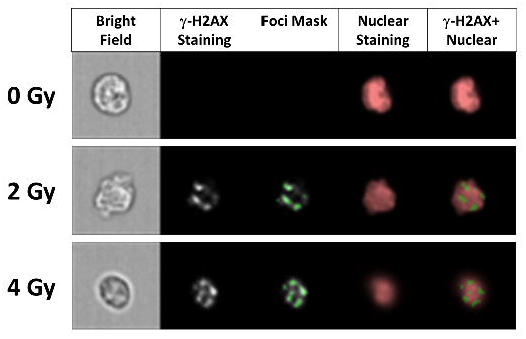
Representative figures of γ-H2AX foci analysis in human blood lymphocytes irradiated cells with γ-rays (0, 2 and 4 Gy), 1 h after irradiation. Images displayed here show cells in bright field, γ-H2AX staining, γ-H2AX foci mask, DRAQ5 nuclear staining and a merged image panel of overlapped γ-H2AX and nuclear staining. The spot counting wizard in the IDEAS® software identified the number of γ-H2AX foci per cell (40X magnification).

### Dose response calibration curve

Figure 5 shows the average dose response for γ-H2AX fluorescence intensity and foci number obtained from 100 µL whole blood samples from four healthy donors, 1 h after 4 Gy exposures. A representative image of non-irradiated human lymphocytes and cells irradiated with 2 Gy and 4 Gy γ rays show non-surprisingly, a higher γ-H2AX fluorescence intensity in the 4 Gy irradiated cells (Figure 5A). The results show a linear increase of γ-H2AX fluorescence intensity (Figure 5B) with increasing radiation dose for the four human donors tested (R^2^=0.9786, p <0.0001). The mean γ-H2AX foci distribution (Figure 5C) indicates that the majority of the control, non-irradiated lymphocyte cells had 0 to 1 γ-H2AX foci, whereas levels ranged from 0 to 8 in the irradiated cells. A small number of cells showed 8-10 differentiable foci after exposure to 4 Gy. The results also show that the linear fit for the mean number of γ-H2AX foci/cell (R^2^=0.8083, p <0.0001, Figure 5D) up to 4 Gy was not as good as using mean γ-H2AX intensity levels.

**Figure 5.**
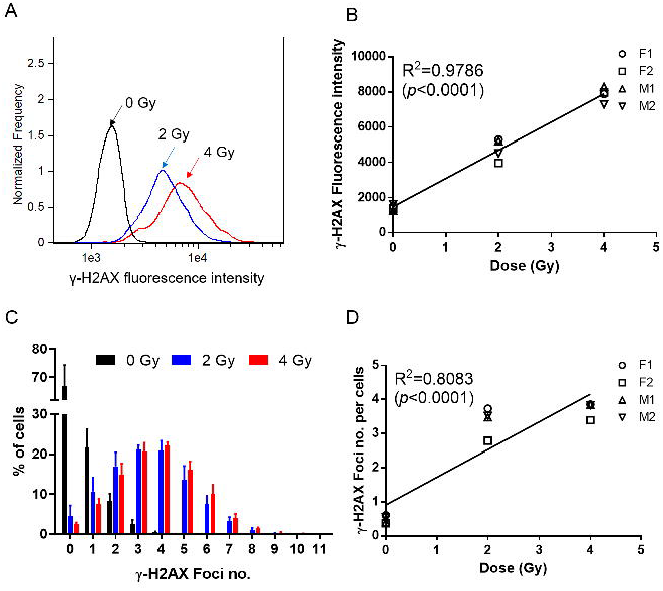
Dose-dependent changes of γ-H2AX in human blood lymphocytes 1 h after exposure with 4 Gy γ rays. (A) Representative distribution of γ-H2AX fluorescence intensity in lymphocytes from a female human donor, F1. (B) Radiation-induced changes in γ-H2AX fluorescence intensity in lymphocytes from 2 female and male donors, F1, F2, M1 and M2. (C) Distribution of cells with different number of γ-H2AX foci in lymphocytes from all donors. Bar indicates mean ± SEM. (D) Radiation-induced changes in γ-H2AX foci number from donors F1, F2, M1 and M2. Each symbol indicates averaged levels of γ-H2AX for each donor and the line represents the mean response.

### Measurement of γ-H2AX yields as a function of time post radiation exposure

Figure 6A shows the time-dependent kinetics for each individual up to 24 h. The results show that radiation-induced γ-H2AX levels rapidly increased within 30 minutes and reached a maximum by ~1 h, after which time there was fast decline by 6 hours, followed by a much slower rate of disappearance up to 24 hours. The kinetics γ-H2AX data are presented using the mean fluorescence intensity measurements because the R^2^ coefficients showed a better fit for this approach, compared to mean foci levels, 0.5 to 24 h post-irradiation (Table 1).

**Table 1.** Dose response of γ-H2AX fluorescence and foci number at different time points

| Time (h) after irradiation | γ-H2AX fluorescence intensity | γ-H2AX foci number per cells |
| --- | --- | --- |
| | $R^2$ ($p$-value) | $R^2$ ($p$-value) |
| 0.5 | 0.9705 (<0.0001) | 0.8295 (<0.0001) |
| 1 | 0.9786 (<0.0001) | 0.8083 (<0.0001) |
| 3 | 0.9533 (<0.0001) | 0.8170 (<0.0001) |
| 6 | 0.8994 (<0.0001) | 0.8540 (<0.0001) |
| 24 | 0.9017 (<0.0001) | 0.7473 (0.0003) |

**Figure 6.**
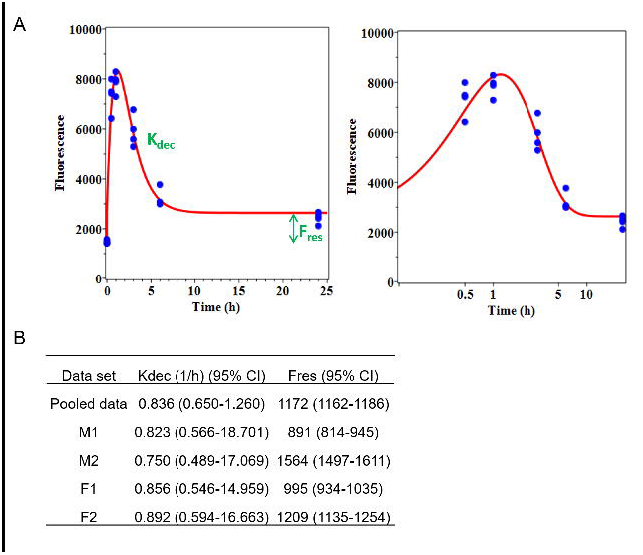
Time-dependent γ-H2AX yields in human blood lymphocytes after 4 Gy irradiation. (A) Experimental data and model fit of γ-H2AX repair kinetics at 0.5, 1, 3, 6 and 24 h after ex vivo irradiation exposure are presented, based on fluorescence intensity; the right panel is the zoomed picture for 0-12 h with a logarithmic time scale which helps to visualize early time points. (B) Each parameter of model fit of γ-H2AX repair kinetics was shown. K_dec_ is the constant for decay of γ-H2AX foci after irradiation. F_res_ is the residual value remaining at long times after irradiation.

Figure 6B shows the data analysis for each individual on γ-H2AX yields as a function of time post radiation exposure. We measured two key parameters to define the γ-H2AX repair kinetics, the rate of decay (K_dec_), and the yield of residual unrepaired breaks (F_res_). Results show the practicality of using this two-parameter approach for quantifying DSB rejoining kinetics.

## Discussion

Since it was first demonstrated by Rogakou, Bonner and colleagues that histone H2AX is rapidly phosphorylated on residue serine 139 in cells when DSBs are introduced into the DNA by ionizing radiation (37), the γ-H2AX assay has been widely used as a sensitive molecular marker of DNA damage and DSB repair capacity in a variety of human tissue and cell types (38, 39). In recent years, the γ-H2AX biomarker has become a powerful tool to monitor DNA DSBs in translational cancer research with the potential to assess the radiosensitivity of prospective radiotherapy patients (5, 40). The goal of the present work was to develop and optimize the γ-H2AX immunocytochemical protocol for high-content screening of double-stranded DNA breaks in finger-stick sized blood samples based on imaging flow cytometry (IFC). The IFC technique allows fast and accurate analysis of γ-H2AX yields in 1000s of cells which would not be possible using conventional manual immunocytochemical protocols.

To assess DSB repair capacity, radiation-induced γ-H2AX yields were measured for dose/time response in in ex-vivo irradiated blood samples taken from four individuals (2 male and female). Measurements of γ-H2AX fluorescence intensity and foci number at specific time points up to 24 hours after exposure with 0, 2 and 4 Gy gamma rays showed linear dose-dependent response and pattern of DNA repair, consistent with previous studies (10, 17, 20, 41). The results highlight that the fluorescence intensity endpoint showed a better dose response compared to foci number given the small difference in foci number between 2 and 4 Gy. The reason for this might be due to the fact that we used the 40X lens for image analysis as opposed to a 60X lens (42, 43). Recently, Parris and colleagues demonstrated the use of higher magnification and application of the focus stacking, called extended depth of field (EDF) increased γ-H2AX foci number and resolution in fibroblast cell line (43).

Quantitative modeling of DNA repair kinetics based on fluorescence intensity showed that the decay constant of γ-H2AX foci after irradiation (K_dec_) was not different among donors tested, whereas residual γ-H2AX fluorescence intensity (F_res_) was apparently higher in M2 and F2 than in the other two donors (M1 and F1), suggesting that M2 and F2 may have more unrepaired DSB 24 h after irradiation (Fig. 6). The differences in DSB repair capacity between the 4 healthy donors tested here, show the potential of our high-throughput γ-H2AX assay to measure DNA repair kinetics on an individual-by-individual basis. Recent work by Kroeber et al showed the capability of the γ-H2AX assay to identify distinct outliers among a large cohort of 136 rectal cancer patients. They identified these patients are most probably radiosensitive and may have the highest risk of suffering radiotherapy-related late sequelae (23). Interestingly, Yin et al recently reported enhanced DNA repair capacity in the peripheral blood mononuclear cells from a small cohort lung cancer patients tended to be associated with a poor response to radiation therapy, implicating a modulation of DNA repair (8).

One benefit of employing IFC technology for γ-H2AX cells analysis is the ability to target specific cell populations as well as eliminate interfering cells or debris which will increase the sensitivity of the assay. In the current study, we measured γ-H2AX yields in focused DNA positive lymphocytes population instead of the total leukocytes. It is known that the sensitivity of lymphocytes and granulocytes to radiation are different whereby γ-H2AX levels in lymphocytes increased in a dose-dependent manner after 0 – 10 Gy γ-ray exposure, but the levels in granulocytes was not affected (36). Further, residual levels of apoptosis in the irradiated samples are a potential confounding factor for the γ-H2AX total fluorescence analysis(44). IFC image analysis using the IDEAS^®^ software allowed us to rapidly detect and eliminate pan-nuclear γ-H2AX stained lymphocytes based on fluorescence intensity and morphology. Pan-nuclear γ-H2AX response has been suggested as a biomarker to distinguish apoptotic cells from DNA damaged cells (45, 46). We showed here that the percentage of pan-nuclear γ-H2AX stained lymphocytes increased over time, up to 24 h after 4 Gy exposure. These observations are consistent with other studies which show the apoptotic response of human lymphocytes upon radiation exposure (47–49).

Another advantage of our IFC-based γ-H2AX assay is reduced assay time and time-to-result. Firstly, our immunolabeling protocol presented here can be completed within 2 hours, eliminating the need of preparation for peripheral blood mononuclear cells which is subjected to Ficoll gradient purification. This involves laborious and time-consuming working steps, which hamper large-scale population studies (50). Additionally, IFC is capable of acquiring cellular imagery at high flow rates, reaching up to 1,000 cells/s, and enabling several different structures within the cell to be analyzed, making it faster than the microscopic system (51). In future studies, we will plan to further automate the IFC γ-H2AX assay system using our own Rapid Automated Biodosimetry Technology (RABiT) platform for automated sample preparation from small volumes of blood (35). We plan also to extend the IFC γ-H2AX assay protocol as a quantitative multiplexed assay to analyze multiple biomarkers on a single cell. Overall, the further development of our IFC-based γ-H2AX system will therefore allow us evaluate DNA damage and DSB repair capacity with increased resolution, sensitivity, accuracy and high-speed image acquisition compared to traditional flow cytometry and traditional microscope immunohistochemical methods (28, 30).

## Conclusions

We have developed a high throughput IFC-based γ-H2AX assay which is a faster and more efficient technique for assessing global DSB repair capacity. These studies could potentially pave the way for new individualized therapy approaches and new large‐scale molecular‐epidemiological studies, with the long‐term goal of predicting individual radiosensitivity and risk of developing adverse effects related to radiotherapy treatment.

## Declarations

### Ethics approval and consent to participate

All procedures involving human participants have been performed in accordance with the Declaration of Helsinki and approved by the Columbia University Medical Center Institutional Review Board (IRB protocol IRB-AAAE-2671). Written informed consent was obtained from all donors.

### Consent for publication

Not applicable

### Availability of data and material

The data that support the findings of this study are available from the corresponding author upon reasonable request.

### Authors’ contributions

YL, HCT, and DJB designed the study. Funding was obtained by DJB. YL, QW and HCT established a protocol for IFC-based γ-H2AX assay. YL and QW performed IFC-based γ-H2AX assay. IS performed modeling of DNA repair kinetics. IS and YL carried out statistical analysis. YL, QW, IS and HCT wrote the manuscript. All authors reviewed and approved the manuscript.

## Acknowledgements

We thank Maria Taveras for blood collection.

## Authors’ information

YL’s current address: Laboratory of Biological Dosimetry, National Radiation Emergency Medical Center, Korea Institute of Radiological and Medical Sciences, 75 Nowon-ro, Nowon-gu, Seoul, Republic of Korea 01812

## References

1. Bau DT, Mau YC, Ding SL, Wu PE, Shen CY. DNA double-strand break repair capacity and risk of breast cancer. Carcinogenesis. 2007;28(8):1726–30.

2. Ralhan R, Kaur J, Kreienberg R, Wiesmüller L. Links between DNA double strand break repair and breast cancer: Accumulating evidence from both familial and nonfamilial cases. Cancer Letters. 2007;248(1):1–17.

3. Parshad R, Sanford KK. Radiation-induced chromatid breaks and deficient DNA repair in cancer predisposition. Critical Reviews in Oncology / Hematology. 2001;37(2):87–96.

4. Rube CE, Grudzenski S, Kuhne M, Dong X, Rief N, Lobrich M, et al. DNA double-strand break repair of blood lymphocytes and normal tissues analysed in a preclinical mouse model: implications for radiosensitivity testing. Clin Cancer Res. 2008;14(20):6546–55.

5. Herschtal A, Martin RF, Leong T, Lobachevsky P, Martin OA. A Bayesian Approach for Prediction of Patient Radiosensitivity. International journal of radiation oncology, biology, physics. 2018;102(3):627–34.

6. Fernet M, Hall J. Genetic biomarkers of therapeutic radiation sensitivity. DNA repair. 2004;3(8-9):1237–43.

7. Chistiakov DA, Voronova NV, Chistiakov PA. Genetic variations in DNA repair genes, radiosensitivity to cancer and susceptibility to acute tissue reactions in radiotherapy-treated cancer patients. Acta oncologica. 2008;47(5):809–24.

8. Yin X, Mason J, Lobachevsky PN, Munforte L, Selbie L, Ball DL, et al. Radiation Therapy Modulates DNA Repair Efficiency in Peripheral Blood Mononuclear Cells of Patients With Non-Small Cell Lung Cancer. International journal of radiation oncology, biology, physics. 2019;103(2):521–31.

9. Valdiglesias V, Giunta S, Fenech M, Neri M, Bonassi S. gammaH2AX as a marker of DNA double strand breaks and genomic instability in human population studies. Mutat Res. 2013;753(1):24–40.

10. Redon CE, Dickey JS, Bonner WM, Sedelnikova OA. gamma-H2AX as a biomarker of DNA damage induced by ionizing radiation in human peripheral blood lymphocytes and artificial skin. Adv Space Res. 2009;43(8):1171–8.

11. Pilch DR, Sedelnikova OA, Redon C, Celeste A, Nussenzweig A, Bonner WM. Characteristics of gamma-H2AX foci at DNA double strand breaks sites. Biochemistry and Cell Biology-Biochimie Et Biologie Cellulaire. 2003;81(3):123–9.

12. Turner HC, Brenner DJ, Chen Y, Bertucci A, Zhang J, Wang H, et al. Adapting the gamma-H2AX assay for automated processing in human lymphocytes. 1. Technological aspects. Radiation research. 2011;175(3):282–90.

13. Redon CE, Nakamura AJ, Sordet O, Dickey JS, Gouliaeva K, Tabb B, et al. gamma-H2AX detection in peripheral blood lymphocytes, splenocytes, bone marrow, xenografts, and skin. Methods Mol Biol. 2011;682:249–70.

14. Rogakou EP, Boon C, Redon C, Bonner WM. Megabase chromatin domains involved in DNA double-strand breaks in vivo. The Journal of cell biology. 1999;146(5):905–16.

15. Sedelnikova OA, Pilch DR, Redon C, Bonner WM. Histone H2AX in DNA damage and repair. Cancer Biol Ther. 2003;2(3):233–5.

16. Beels L, Werbrouck J, Thierens H. Dose response and repair kinetics of gamma-H2AX foci induced by in vitro irradiation of whole blood and T-lymphocytes with X- and gamma-radiation. Int J Radiat Biol. 2010;86(9):760–8.

17. Rothkamm K, Horn S. gamma-H2AX as protein biomarker for radiation exposure. Annali dell’Istituto superiore di sanita. 2009;45(3):265–71.

18. Bhogal N, Kaspler P, Jalali F, Hyrien O, Chen R, Hill RP, et al. Late residual gamma-H2AX foci in murine skin are dose responsive and predict radiosensitivity in vivo. Radiat Res. 2010;173(1):1–9.

19. Slyskova J, Naccarati A, Polakova V, Pardini B, Vodickova L, Stetina R, et al. DNA damage and nucleotide excision repair capacity in healthy individuals. Environmental and molecular mutagenesis. 2011;52(7):511–7.

20. Sharma PM, Ponnaiya B, Taveras M, Shuryak I, Turner H, Brenner DJ. High throughput measurement of gammaH2AX DSB repair kinetics in a healthy human population. PloS one. 2015;10(3):e0121083.

21. Mumbrekar KD, Goutham HV, Vadhiraja BM, Bola Sadashiva SR. Polymorphisms in double strand break repair related genes influence radiosensitivity phenotype in lymphocytes from healthy individuals. DNA Repair (Amst). 2016;40:27–34.

22. Lobachevsky P, Leong T, Daly P, Smith J, Best N, Tomaszewski J, et al. Compromized DNA repair as a basis for identification of cancer radiotherapy patients with extreme radiosensitivity. Cancer Lett. 2016;383(2):212–9.

23. Kroeber J, Wenger B, Schwegler M, Daniel C, Schmidt M, Djuzenova CS, et al. Distinct increased outliers among 136 rectal cancer patients assessed by gammaH2AX. Radiat Oncol. 2015;10:36.

24. Nagel ZD, Engelward BP, Brenner DJ, Begley TJ, Sobol RW, Bielas JH, et al. Towards precision prevention: Technologies for identifying healthy individuals with high risk of disease. Mutation Research/Fundamental and Molecular Mechanisms of Mutagenesis. 2017;800–802:14–28.

25. Basiji DA, Ortyn WE, Liang L, Venkatachalam V, Morrissey P. Cellular image analysis and imaging by flow cytometry. Clin Lab Med. 2007;27(3):653-+.

26. Basiji D, O’Gorman MR. Imaging flow cytometry. J Immunol Methods. 2015;423:1–2.

27. Basiji DA. Principles of Amnis Imaging Flow Cytometry. Methods in molecular biology. 2016;1389:13–21.

28. Vorobjev IA, Barteneva NS. Quantitative Functional Morphology by Imaging Flow Cytometry. Methods Mol Biol. 2016;1389:3–11.

29. Vorobjev IA, Barteneva NS. Temporal Heterogeneity in Apoptosis Determined by Imaging Flow Cytometry. Methods Mol Biol. 2016;1389:221–33.

30. Han Y, Gu Y, Zhang AC, Lo YH. Review: imaging technologies for flow cytometry. Lab Chip. 2016;16(24):4639–47.

31. Doan M, Vorobjev I, Rees P, Filby A, Wolkenhauer O, Goldfeld AE, et al. Diagnostic Potential of Imaging Flow Cytometry. Trends Biotechnol. 2018;36(7):649–52.

32. Barteneva NS, Fasler-Kan E, Vorobjev IA. Imaging flow cytometry: coping with heterogeneity in biological systems. J Histochem Cytochem. 2012;60(10):723–33.

33. Dias AM, Almeida CR, Reis CA, Pinho SS. Studying T Cells N-Glycosylation by Imaging Flow Cytometry. Methods Mol Biol. 2016;1389:167–76.

34. Pischel D, Buchbinder JH, Sundmacher K, Lavrik IN, Flassig RJ. A guide to automated apoptosis detection: How to make sense of imaging flow cytometry data. PLoS One. 2018;13(5):e0197208.

35. Wang Q, Rodrigues MA, Repin M, Pampou S, Beaton-Green LA, Perrier J, et al. Automated Triage Radiation Biodosimetry: Integrating Imaging Flow Cytometry with High-Throughput Robotics to Perform the Cytokinesis-Block Micronucleus Assay. Radiat Res. 2019;191(4):342–51.

36. Wang Z, Hu H, Hu M, Zhang X, Wang Q, Qiao Y, et al. Ratio of γ-H2AX level in lymphocytes to that in granulocytes detected using flow cytometry as a potential biodosimeter for radiation exposure. Radiat Environ Biophys. 2014;53(2):283–90.

37. Rogakou EP, Pilch DR, Orr AH, Ivanova VS, Bonner WM. DNA double-stranded breaks induce histone H2AX phosphorylation on serine 139. J Biol Chem. 1998;273(10):5858–68.

38. Mah LJ, El-Osta A, Karagiannis TC. gammaH2AX: a sensitive molecular marker of DNA damage and repair. Leukemia. 2010;24(4):679–86.

39. Leatherbarrow EL, Harper JV, Cucinotta FA, O’Neill P. Induction and quantification of gamma-H2AX foci following low and high LET-irradiation. Int J Radiat Biol. 2006;82(2):111–8.

40. Ivashkevich A, Redon CE, Nakamura AJ, Martin RF, Martin OA. Use of the gamma-H2AX assay to monitor DNA damage and repair in translational cancer research. Cancer Lett. 2012;327(1-2):123–33.

41. Wilkins RC, Rodrigues MA, Beaton-Green LA. The Application of Imaging Flow Cytometry to High-Throughput Biodosimetry. Genome Integr. 2017;8:7.

42. Durdik M, Kosik P, Gursky J, Vokalova L, Markova E, Belyaev I. Imaging flow cytometry as a sensitive tool to detect low-dose-induced DNA damage by analyzing 53BP1 and gammaH2AX foci in human lymphocytes. Cytometry A. 2015;87(12):1070–8.

43. Parris CN, Adam Zahir S, Al-Ali H, Bourton EC, Plowman C, Plowman PN. Enhanced gamma-H2AX DNA damage foci detection using multimagnification and extended depth of field in imaging flow cytometry. Cytometry A. 2015;87(8):717–23.

44. Turner HC, Shuryak I, Taveras M, Bertucci A, Perrier JR, Chen C, et al. Effect of dose rate on residual gamma-H2AX levels and frequency of micronuclei in X-irradiated mouse lymphocytes. Radiat Res. 2015;183(3):315–24.

45. Ding D, Zhang Y, Wang J, Zhang X, Gao Y, Yin L, et al. Induction and inhibition of the pan-nuclear gamma-H2AX response in resting human peripheral blood lymphocytes after X-ray irradiation. Cell Death Discov. 2016;2:16011.

46. Solier S, Pommier Y. The apoptotic ring: a novel entity with phosphorylated histones H2AX and H2B and activated DNA damage response kinases. Cell Cycle. 2009;8(12):1853–9.

47. Belloni P, Meschini R, Czene S, Harms-Ringdahl M, Palitti F. Studies on radiation-induced apoptosis in G0 human lymphocytes. Int J Radiat Biol. 2005;81(8):587–99.

48. Boreham DR, Dolling JA, Maves SR, Siwarungsun N, Mitchel RE. Dose-rate effects for apoptosis and micronucleus formation in gamma-irradiated human lymphocytes. Radiat Res. 2000;153(5 Pt 1):579–86.

49. Payne CM, Bjore CG, Jr., Schultz DA. Change in the frequency of apoptosis after low- and high-dose X-irradiation of human lymphocytes. J Leukoc Biol. 1992;52(4):433–40.

50. Ismail IH, Wadhra TI, Hammarsten O. An optimized method for detecting gamma-H2AX in blood cells reveals a significant interindividual variation in the gamma-H2AX response among humans. Nucleic Acids Res. 2007;35(5):e36.

51. Heylmann D, Kaina B. The gammaH2AX DNA damage assay from a drop of blood. Sci Rep. 2016;6:22682.

